# Modeling of individual neurophysiological brain connectivity

**DOI:** 10.1101/2022.03.02.482608

**Authors:** S.D. Kulik, L. Douw, E. van Dellen, M.D. Steenwijk, J.J.G. Geurts, C.J. Stam, A. Hillebrand, M.M. Schoonheim, P. Tewarie

## Abstract

**Introduction:** Computational models are often used to assess how functional connectivity (FC) patterns emerge from neuronal population dynamics and anatomical connections in the brain. However, group averaged data is often used in this context and it remains unclear whether individual predictions of FC patterns using this approach can be made. Here, we assess the value of using individualized structural data for simulation of individual whole-brain FC.

**Methods:** The Jansen and Rit neural mass model was employed, where masses were coupled using individual structural connectivity (SC) obtained from diffusion weighted imaging. Simulated FC was correlated to individual magnetoencephalography-derived empirical FC. FC was estimated using both phase-based (phase lag index (PLI), phase locking value (PLV)) and amplitude-based (amplitude envelope correlation (AEC)) metrics to analyze the goodness-of-fit of different metrics for individual predictions. Prediction of individual FC was compared against the prediction of group averaged FC. We further tested whether SC of a different participant could equally well predict a participants FC pattern.

**Results:** The AEC provided a significantly better match between individually simulated and empirical FC than phase-based metrics. Simulations with individual SC provided higher correlations between simulated and empirical FC compared to using the group-averaged SC. However, using SC from other participants resulted in similar correlations between simulated and empirical FC compared to using participants own SC.

**Discussion:** This work underlines the added value of FC simulations based on individual instead of group-averaged SC, and could aid in a better understanding of mechanisms underlying individual functional network trajectories in neurological disease.

**Impact statement:** In this work, we investigated how well individual empirical functional connectivity can be simulated using the individual’s structural connectivity matrix combined with neural mass modeling. Our research highlights the potential added value of using individual simulations of functional connectivity, and could aid in a better understanding of mechanisms underlying individual functional network trajectories in neurological disease. Moreover, individualized prediction of disease trajectories could enhance patient care and may provide better treatment options.

## Introduction

The brain is a complex network of brain regions that display interregional communication, i.e. so called functional connectivity (FC). FC is defined by statistical interdependencies between time-series of brain activity (Friston 2011). In case of neurophysiological data, FC can be estimated from either the phase or amplitude of neuronal oscillations (Siegel et al 2012, Siems & Siegel 2020). Disruption of the FC patterns are known to be clinically relevant in neurological (Stam 2014) and psychiatric disorders (Hallett et al 2020). Computational models are often used to gain insight into mechanisms that result in disrupted patterns of FC. Using this approach, the impact of pathology at the neuronal population level or at the level of structural connections (SC) on FC can be assessed and used to make predictions of empirical FC patterns. Especially individualized prediction of disease trajectories (Douw et al 2019) are important in this context. However, so far mainly group averaged SC and FC have been used, and it remains an open question whether individual predictions of FC are feasible, even in healthy conditions.

Computational modeling of brain activity and FC can be approached using so-called neural mass modelling (Deco et al 2008). Neural mass models assume a mean ensemble activity of neurons that reduces the number of dimensions and allows multiple interacting local populations (Breakspear 2017). A neural mass corresponds to activity within a brain region and masses can be coupled using empirically measured structural connections, resulting in whole-brain network simulations. A well-known model that is known to generate physiologically accurate brain activity (Aburn et al 2012) was developed by (Lopes da Silva et al 1974) and further improved by Jansen and Rit (Jansen & Rit 1995). The Jansen and Rit model is able to produce oscillatory activity in the alpha band, i.e. the dominant rhythm in resting-state neurophysiological data. Usage of this model can be justified by the fact that its dynamical properties have been thoroughly investigated and are well understood (Grimbert & Faugeras 2006, Spiegler et al 2011).

So far, computational modeling of empirical neurophysiological connectivity is mainly based on group-averaged SC as input to neural mass models (Abeysuriya et al 2018, Cabral et al 2014, Deco et al 2017, Hadida et al 2018, Moon et al 2015, O’Neill et al 2018, Tewarie et al 2019a, Tewarie et al 2014). One previous study on structure-function relationships compared individually simulated and empirically derived FC, based on electroencephalography (EEG) data (Finger et al 2016). This study showed moderate to strong correlations between individually simulated and empirical FC by using a simple autoregressive model. FC was calculated with different phase-based FC metrics. Finger and colleagues tested the specificity of using individual SC by correlating individually simulated FC with either the corresponding empirical FC matrices, or with empirical FC matrices of other participants, and found no significant differences between the two approaches. This finding could be supported by a recent functional magnetic resonance imaging (MRI) study (Zimmermann et al 2019) where it was found that the correspondence between empirical SC and FC in many participants was limited due to the small variability between participants in SC compared with the larger variability in FC, perhaps indicating that structural data is not specific enough to simulate FC accurately. Despite the relevance of previous work (Finger et al 2016), we argue that the feasibility of individual predictions of FC should be re-tested in an independent dataset and should be tested using both amplitude- and phase-based metrics for FC, as recent work suggest that both phase and amplitude could encode complementary information (Siems & Siegel 2020). However, this observation has not been reproduced in an independent dataset. In addition, we will extend previous work by including more participants, making use of magnetoencephalography (MEG) instead of EEG data and applying different FC metrics.

In the current work, we investigated how well individual empirical FC can be approximated by simulating an estimate of FC based on an individual’s own SC. We analyzed both amplitude and phase-based metrics in this context, calculated from MEG data. In order to put our results into perspective, we compared our results of individual simulations with FC approximations based on group-averaged SC and individual predictions based on non-matched empirical SC.

## Methods

### Participants

Forty healthy participants (37.5% men, age 50.7 ± 6.1 years) from the Amsterdam multiple sclerosis cohort were included (Eijlers et al 2018). We only included participants who underwent both diffusion MRI (dMRI) and magnetoencephalography recordings. Approval was obtained from the institutional ethics review board of the VU University Medical Center, and participants gave written informed consent prior to participation.

### Empirical structural data: diffusion MRI

Individually weighted dMRI matrices were obtained to describe the structural connectivity between the neural masses. dMRI matrices were calculated with probabilistic tractography as described previously (Meijer et al 2020). In short, participants were scanned on a 3 T scanner (GE signa HDxt), using an eight-channel phased-array head-coil. For volumetric measurements, a 3D T1-weighted inversion-prepared fast spoiled gradient recall sequence (FSPGR, repetition time 7.8 ms, echo time 3 ms, inversion time 450 ms, flip angle 12°, sagittal 1.0 mm sections, 0.94 × 0.94 mm^2^ in-plane resolution) was taken into account. A diffusion weighted imaging sequence (dMRI) was applied covering the entire brain using five volumes without directional weighting (i.e. b=0 s/mm^2^) and 30 volumes with non-collinear diffusion gradients (echo planar imaging (EPI), b=1000 s/mm^2^, repetition time 13000 ms, echo time 91 ms, flip angle 90°, 2.4 mm contiguous axial slices, 2 × 2 mm^2^ in-plane resolution). Subsequently, the FMRIB Diffusion Toolbox (FDT; part of FSL 5) was performed using eddy current distortion correction. Next, using the fiber orientation distribution (FOD), probabilistic tractography was applied using MRtrix 3.0 (Tournier et al 2012). In this model, *N* streamlines are reconstructed by randomly putting seeds in white matter and using constrained spherical deconvolution to estimate the local FOD (Tournier et al 2007). The 30 non-collinear diffusion directions in the data were adjusted by restricting the maximum spherical harmonic order (lmax) to 6. Then, for each participant, a random seeding of 100 million fibers within the brain mask was applied to perform whole-brain probabilistic tractography. Probabilistic tractography was applied because it is frequently used due to its low sensitivity for false-positives (Maier-Hein et al 2017).

Cortical grey matter regions were defined by processing the 3D T1-weighted image of each participant with the FreeSurfer 5.3 pipeline. The automated anatomical labeling (AAL) atlas (Tzourio-Mazoyer et al 2002) was used to define 78 cortical regions (Gong et al 2009) on the native cortical surface. Structural networks were constructed by considering regions as nodes and the number of fibers between pairs of nodes as links. We performed normalization of elements in the SC matrices. For each individual SC matrix, link weights that exceeded 1.5*IQR (interquartile range) above the third quartile (Q3 + 1.5*IQR) were set to that value, to make sure that very high values would not disproportionally influence the simulations. Subsequently, the weighted structural connectivity matrices were rescaled to the range [0 1].

### Empirical functional data: magnetoencephalography

Acquisition and pre-processing of the MEG data was performed as described previously (Derks et al 2018). In short, eyes-closed, resting-state measurements of 5 minutes were used. Measurements were performed in a magnetically shielded room (Vacuum Schmelze GmbH, Hanua, Germany) with a 306-channel whole-head MEG system (Elekta Neuromag Oy, Helsinki, Finland). Data were sampled at 1250 Hz, and a high-pass filter (0.1 Hz) and anti-aliasing filter (410 Hz) were employed online. The extended Signal Space Separation method (xSSS) (van Klink et al 2017) was applied, after which a maximum of 12 malfunctioning channels were excluded during visual inspection (SK, LD). Artefact removal was performed offline with the temporal extension of SSS in MaxFilter software (Elekta Neuromag Oy, version 2.2.15) (Taulu & Simola 2006). The head position relative to the MEG sensors was recorded continuously with the signals from four or five head-localization coils. The head-localization coil positions and outline of the participants scalp were digitized using a 3D digitizer (3Space Fastrak, Polhemus, Colchester, VT, USA). Each participant’s scalp surface was co-registered to their structural MRI using a surface-matching procedure. Subsequently, the co-registered MRI was spatially normalized to a template MRI. Centroid voxels (Hillebrand et al 2016) of the 78 cortical regions of the AAL atlas, the same as was used for the SC, were selected for further analyses after inverse transformation to the participant’s co-registered MRI. A single best fitting sphere was fitted to the outline of the scalp as obtained from the co-registered MRI and used as a volume conductor model for the beamformer approach (Hillebrand & Barnes 2005, Hillebrand et al 2005). An atlas-based scalar beamformer implementation (Elekta Neuromag Oy, version 2.1.28), similar to Synthetic Aperture Magnetometry (Robinson & Vrba 1999), was applied to project MEG data from sensor level to source space (Hillebrand et al 2012). The beamformer weights were based on the data covariance matrix and the forward solution (lead field) of a dipolar source at the voxel location. Orientation of the sources was estimated based on singular value decomposition (Sekihara et al 2006). The broadband (0.5-48 Hz) time-series of the 78 centroids were projected through the normalized (Cheyne et al 2007) broadband beamformer weights for each target voxel (i.e. centroid voxel). From these time-series, for each participant, the maximum amount of artefact free data, i.e. 26 consecutive epochs of 6.55 seconds (8192 samples), were analyzed (Liuzzi et al 2017). Time-series were digitally band-pass filtered in the alpha band (8-13 Hz) using a fast Fourier transform, after which all bins outside the passbands were set to zero, and an inverse Fourier transform was performed (implemented using in house script in Matlab (version 2018b, Mathworks, Natick, MA, USA)). Subsequently, FC was calculated using different FC metrics (see paragraph *‘Simulated and empirical functional connectivity’*). All the analyses in the current work were performed in Matlab using in house scripts (see https://github.com/multinetlab-amsterdam/projects/tree/master/modelling_paper_2021).

### Simulated functional data: network of neural masses

We considered a network of coupled neural masses with network size *N* = 78. Each node (neural mass) corresponded to a cortical region of the AAL atlas. Link weights (number of streamlines) were derived from an individual’s weighted SC matrix. We used the Jansen and Rit model as described in (Grimbert & Faugeras 2006) to model a single neural mass. This model allows for simulation of fluctuations in the synaptic membrane potential of a neuronal population (Jansen & Rit 1995). Each mass consists of three populations (pyramidal population, and excitatory and inhibitory neuronal populations) (see Figure 1A). The Jansen and Rit model is optimized to generate alpha oscillations. In short, each neuronal population is described by a second-order ordinary differential equation that models modulations in the mean membrane potential due to the mean incoming firing rate from the same population and from other populations in the neural mass. Incoming mean firing rates are obtained by a nonlinear sigmoid function that transforms the mean membrane potential to a mean firing rate (Jansen & Rit 1995). Uncorrelated Gaussian noise was fed to the pyramidal population only. The three interconnected neuronal populations were connected using the coupling values (C1, C2, C3, C4) (Figure 1 and Table 1). These values represent the average number of synaptic connections between each population. Connectivity between the neural masses was implemented exactly the same as in (Forrester et al 2020) and the same parameters were used as in (Grimbert & Faugeras 2006). A fourth order stochastic Runge-Kutta method (Hansen & Penland 2006) was used to numerically solve the coupled differential equations of the model.

**Figure 1.**
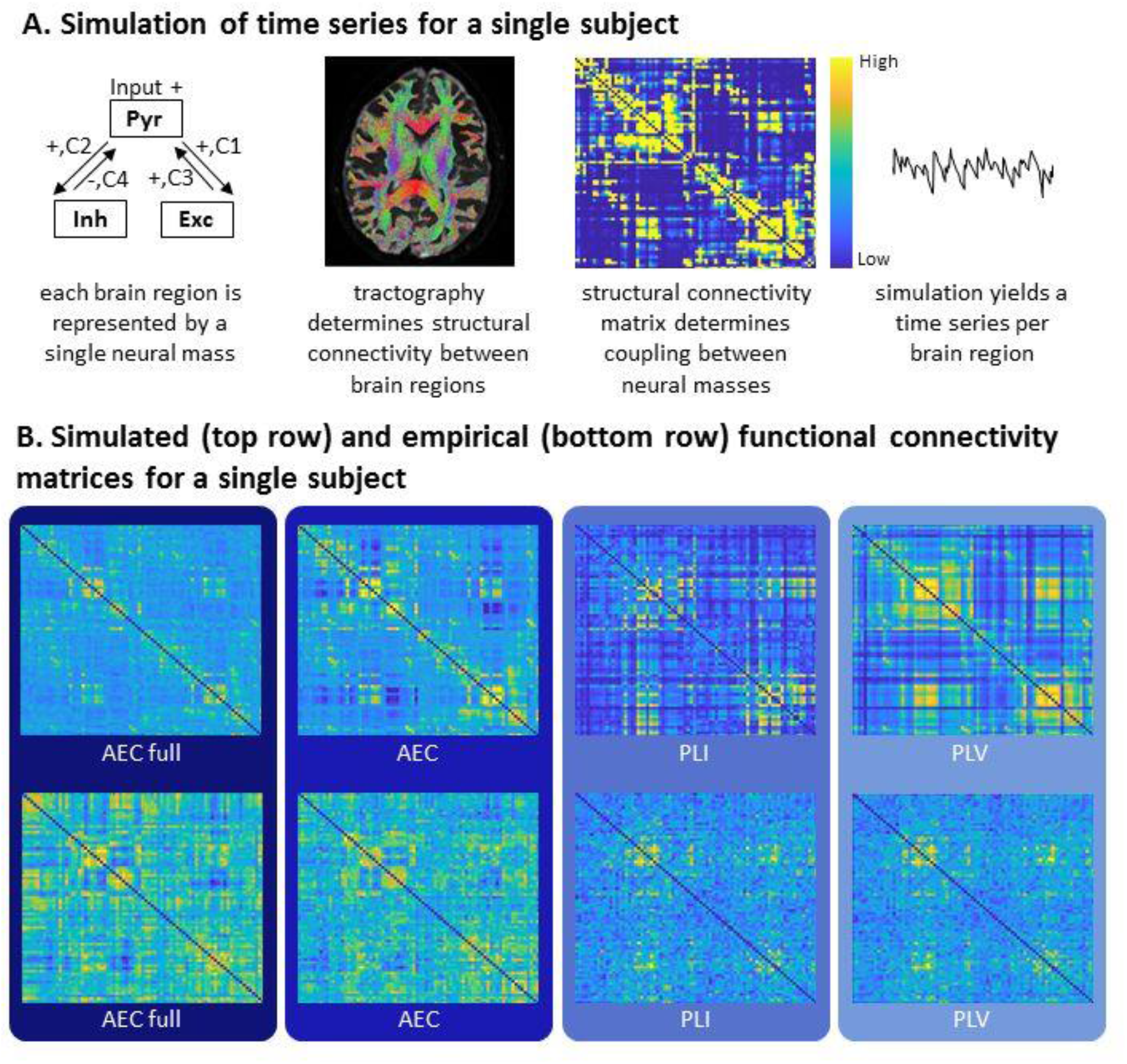
Overview of the applied methods. A: left: overview of the Jansen and Rit model reflecting the connections between the pyramidal (Pyr), inhibitory (Inh) and excitatory (Exc) populations. Individual weighted structural connectivity, computed by probabilistic tractography using MRTrix, was used as input to the Jansen and Rit model to connect the neural masses. Each neural mass, reflecting a brain region, produces MEG-like time series. B: Exemplar simulated and empirical weighted functional connectivity matrices for one participant. Cold colors represent low connectivity and warmer colors represent high connectivity (this also applies to the structural connectivity matrix). For both simulated and empirical data, FC was estimated between all pairs of regions using different FC metrics (AEC: amplitude envelope correlation, AEC full: AEC calculated over the full time-series (epochs concatenated), AEC: calculated over epochs, PLI: phase lag index, calculated over epochs, PLV: phase locking value, calculated over epochs). AEC full, AEC and PLV were corrected for signal leakage in the empirical data, not in the simulated data. PLI inherently corrects for signal leakage and therefore corrects in both empirical and simulated data. For each participant and per connectivity metric, a correlation between the simulated and empirical FC was performed.

**Table 1.** Parameters and values included in the model (based on (Grimbert & Faugeras 2006))

| Parameter | Meaning | Value |
| --- | --- | --- |
| **C1, C2, C3, C4** | Average number of synapses between populations | 135 * [1 0.8 0.25 0.25] |
| **Beta_E** | Timescale for excitatory population | 100 ms |
| **Beta_I** | Timescale for inhibitory population | 50 ms |
| **A** | Average excitatory synaptic gain | 3.25 |
| **B** | Average inhibitory synaptic gain | 22 |
| **nu** | Threshold of sigmoid | 5 s<sup>-1</sup> |
| **r** | Slope of sigmoid | 0.56 mV<sup>-1</sup> |
| **theta** | Amplitude of sigmoid | 6 mV |
| **Conduction velocity** |  | 10 m/s |
| **Fs** | Sample frequency | 1250 Hz |
| **h** | Integration time step | 0.0001 |
| **T** | Observation time | 20 s |
| **P** | External input to each of the neural masses | 150 |
| **Coupling** | Coupling between the neural masses | [0.1:0.012:0.292] |

Each neural mass receives external input (P) that corresponds to external sources or activity from neighboring populations (Ableidinger et al 2017). The external input was set to P=150 for all neural masses. For the global coupling parameter, which determines the coupling between all neural masses, we used the interval [0.1, 0.292], with a discrete step size of 0.012. As explained in more detail later, this range was used to scan the parameter space in order to obtain the coupling value for every individual that optimized the goodness of fit between simulated and empirical FC matrices. We included distance dependent delays between nodes based on the Euclidian distance between centroids in the AAL atlas divided by the conduction velocity. See Table 1 for an overview of all model parameters. We ran the model for each global coupling value to generate time-series of neuronal activity. For each run, the time-series were band-pass filtered in the alpha band (8-13 Hz) in the same way as for empirical data, and FC was calculated using different FC metrics (see paragraph *‘Simulated and empirical functional connectivity’*). In order to obtain robust results and in order to minimize the stochastic effect of the model’s stochastic differential equations, the model was ran 20 times, and subsequently FC values were averaged over the 20 runs.

### Simulated and empirical functional connectivity

Three FC metrics were calculated that capture either amplitude-based connectivity or phase-based connectivity: the amplitude envelope correlation (AEC) (Brookes et al 2011, Bruns et al 2000, Hipp et al 2012), the Phase Lag Index (PLI) (Stam et al 2007) and the Phase Locking Value (PLV) (Lachaux et al 1999). The AEC quantifies amplitude-based connectivity between two time-series, whereas the PLI and PLV are both metrics of phase synchronization. The main difference between the latter two metrics is that the PLI inherently is insensitive to zero lag phase differences and thereby reduces the effect of primary signal leakage. Prior to FC estimation, we first band-pass filtered the data in the alpha band (8-13 Hz) followed by correction for signal leakage. More specifically, we applied pairwise orthogonalisation in order to correct for signal leakage only in empirical data and only for metrics that are inherently sensitive to signal leakage (AEC and PLV). To calculate the AEC, the amplitude envelopes were obtained from the analytical signal after a Hilbert transformation of the band-pass filtered orthogonalised time-series, and the correlations between the amplitude envelopes of pairs of time-series were computed. For the empirical data, the AEC was calculated in two different ways: 1) AEC: the data were divided into epochs (6.55 seconds), and AEC computed for every epoch. The AEC was subsequently averaged over epochs; 2) AEC full: AEC was computed for the entire time-series, after concatenating all epochs. To calculate the PLI and the PLV, the instantaneous phases were obtained from the same analytical signal after the Hilbert transformation. The PLI and PLV were both calculated for every epoch (6.55 seconds) and subsequently averaged over epochs. For the simulated data, for each FC metric, the FC matrices were averaged over the 20 runs per coupling value.

### Similarity between simulated and participant-specific empirical functional connectivity using individual structural connectivity

We computed a Spearman rank correlation (*ρ*) between simulated and empirical FC matrices for every global coupling value to quantify the match between simulated and individual empirical FC. To do this, the upper triangular part of the matrices were vectorised and subsequently correlated between simulated and empirical FC. Spearman correlations were applied since the distribution of FC values for most metrics was typically-non-Gaussian. For all statistical tests performed, values of *p* < 0.05 were considered to be significant. Simulations were performed with the individual SC matrix as input to the neural mass models. The highest Spearman correlation within the coupling range [0.1, 0.292] was considered to be the best fit with the empirical FC, further referred to as the maximum correlation per participant, and calculated per FC metric. If the coupling value corresponding to the maximum correlation was at the end of the coupling range (i.e. coupling = 0.292), we extended the coupling range to 0.4, with a step size of 0.012, to test whether that coupling range would result in higher correlation values for that individual. Subsequently, the maximum correlation for the range [0.1, 0.4] was determined.

A Wilcoxon signed rank test was subsequently performed to compare the maximum correlations between FC metrics. The FC metric that resulted in the highest maximum correlations at the group level was selected for further analyses. Differences between coupling values corresponding to the maximum correlations for the different FC metrics were tested with Friedman’s test.

### Similarity between simulated and participant-specific empirical functional connectivity using group-averaged structural connectivity

We subsequently tested whether the individual SC as input to the model outperformed simulations based on the group-averaged SC. We therefore used the average SC as input to the model and correlated the resulting simulated FC for the range of coupling values, with the individual empirical FC, using a Spearman correlation. As reference, we also predicted group-averaged FC based on simulations with the group-averaged SC as input. The group-averaged SC and FC matrices were obtained by averaging SC and FC matrices across all participants, respectively. Next, in the group-averaged weighted SC, outliers were removed and normalization of the matrix was applied as described in section *Empirical structural data: diffusion MRI’*. All subsequent steps to calculate the match between simulated and empirical FC were as described in section *‘Similarity between simulated and participant-specific empirical functional connectivity using individual structural connectivity’*.

### Simulated versus empirical functional connectivity in matched versus non-matched participants

In a subsequent analysis, we tested whether the predictions of individual empirical FC based on participants’ own SC matrix were specific. We tested the null hypothesis that prediction of empirical FC for a given participant based on simulated FC with the SC of another participant as input to the simulations would lead to an equally well prediction. To this end, we correlated the individually simulated FC matrices to empirical FC matrices from other participants. We then compared the Spearman correlations between simulated and empirical FC for matched versus non-matched data. To test whether participant’s own maximum correlation (matched data) was higher compared to the correlations obtained with all other participants’ empirical data (non-matched data), these correlations were ranked per participant. Subsequently, if the participant’s own maximum correlation would fall within the highest 97.5% of this ranking, it was considered to be significantly higher compared to the correlations to other participants.

For all previously described analyses, no corrections for multiple comparisons were performed.

## Results

Exemplar time-series and power spectrum of simulated data for one participant are shown in Supplementary Figure S1. Examples of simulated and empirical FC matrices of the same participant are shown in Figure 1B.

### Similarity between individually simulated and empirical functional connectivity

The similarity between the individually simulated and individual empirical FC was calculated for each of the FC metrics for the range of coupling values. The resulting individual maximum correlation values between simulated and empirical FC are shown in Figure 2 for each FC metric. The median of the maximum correlations for each FC metric were: AEC full 0.19, AEC 0.19, PLI 0.10, PLV 0.14. All of these maximum correlations between simulated and empirical FC for the AEC full and AEC were statistically significant (for all participants with AEC full *p* < 0.001, for all participants with AEC: p < 0.01). For the PLI and PLV, correlations between the simulated and empirical FC were statistically significant for most participants (PLI: *p* < 0.005, PLV: *p* < 0.01), except for three (PLI) and two (PLV) participants. The coupling values corresponding to the maximum correlation between simulated and empirical FC for each participant and each FC metric are displayed in Figure 3 and Supplementary table S1. Coupling values corresponding to the maximum correlations did not differ between metrics (χ^2^ = 5.09, *p* = 0.17).

**Figure 2:**
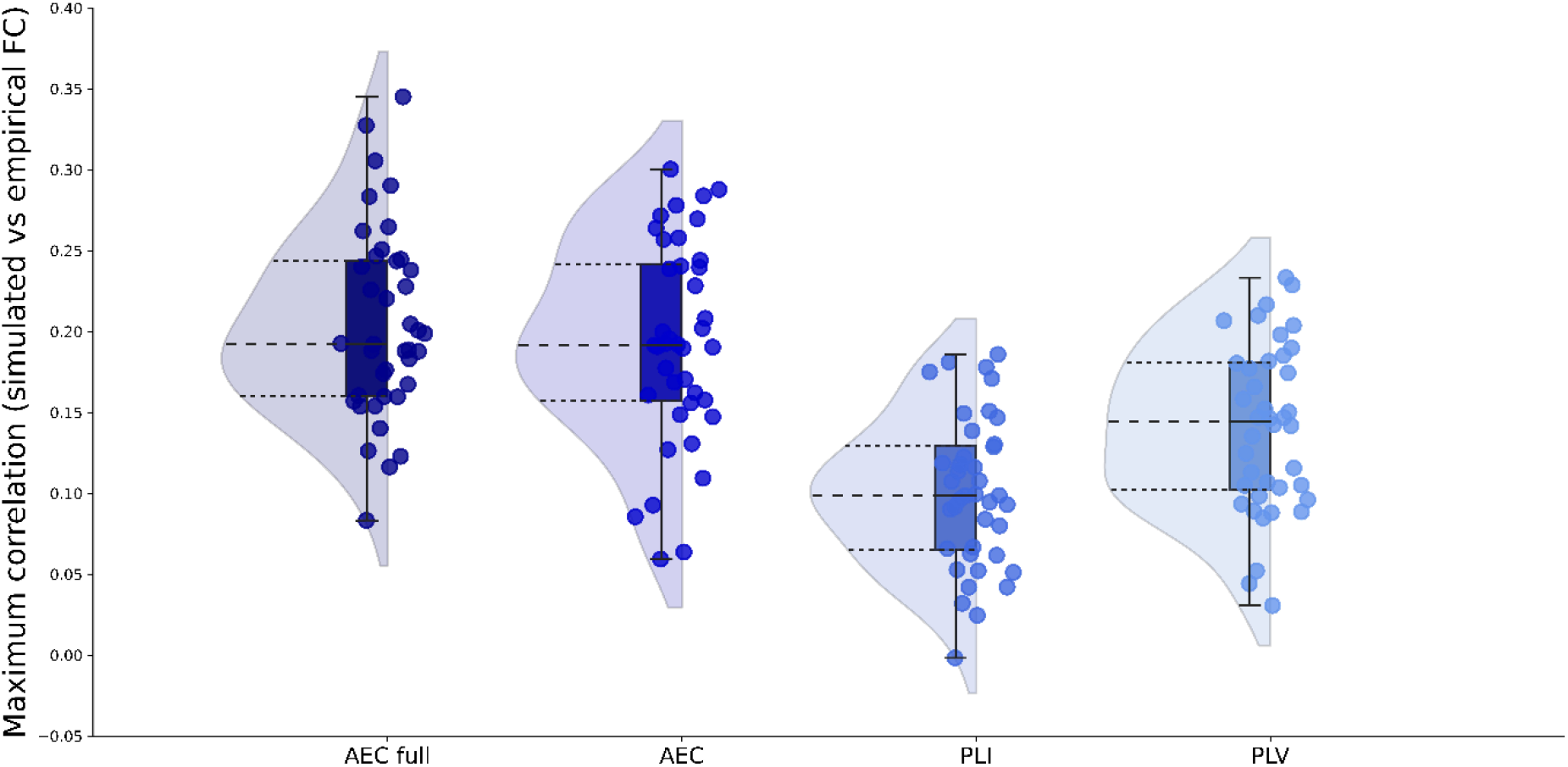
Maximum correlations between simulated and empirical FC. Raincloud figures showing the maximum correlations between simulated and empirical FC for each FC metric. Both amplitude and phase based FC metrics were included: amplitude envelope correlation (AEC); AEC full refers to AEC computed over the full time-series, phase lag index (PLI), phase locking value (PLV).

**Figure 3.**
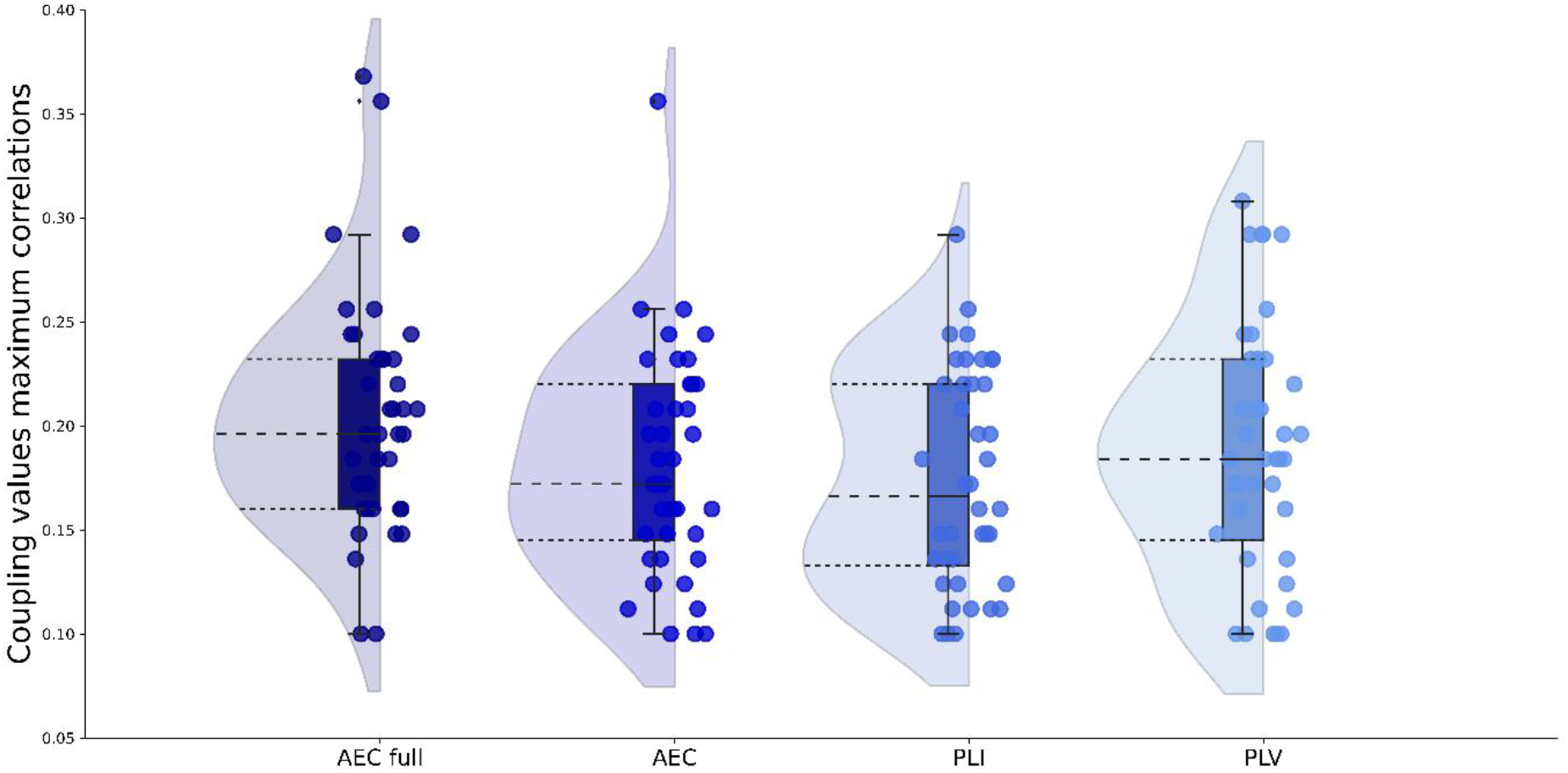
Optimal global coupling values for all FC metrics. Optimized global coupling values between neural masses as determined by the maximum correlation between simulated and empirical FC for each FC metric. Abbreviations: AEC: amplitude envelope correlation (AEC), AEC full: AEC computed over the full time-series, PLI: phase lag index, PLV: phase locking value.

We compared individual maximum correlations between FC metrics. There was no significant difference between the AEC full and the AEC (W = 505, *p* = 0.20). AEC full showed significantly higher maximum correlations than the PLI (W = 804, *p* < 0.001), and the PLV (W = 722, *p* < 0.001). The AEC also showed significantly higher maximum correlations compared to both the PLI (W = 787, *p* < 0.001) and the PLV (W = 699, *p* < 0.001). Finally, the PLI performed significantly worse than the PLV (W = 28, *p* < 0.001) in terms of maximum correlations between simulated and empirical FC at the individual level. Since the use of the AEC full and AEC resulted in significant better predictions of individual empirical FC, we continued using only these metrics for further analyses.

Additionally, we analyzed the similarity between the strongest connections of the individually simulated and empirical data. A detailed description of this analyses can be found in the Supplementary Information. For the AEC, maximal correlations between the strongest connections of simulated and empirical data showed to be significantly higher compared to the maximal correlations when the full matrices were taken into account (W=205, *p* = 0.006, see Figure S2).

### Similarity between simulated functional connectivity and empirical function connectivity using group-averaged structural connectivity

We next predicted individual empirical FC (AEC full and AEC) based on simulations with the group-averaged SC as input. Results show a median of the maximum correlations of 0.19 for both the AEC full and AEC (see Figure 4). There was no significant difference between the maximum correlations for these two FC metrics (W = 462, *p* = 0.5). The match between simulated and individual empirical FC was better for simulations with the individual SC as input compared to simulations with the group-averaged SC as input, for both the AEC full (W = 185, *p* = 0.003) and AEC (W = 200, *p* = 0.005).

**Figure 4:**
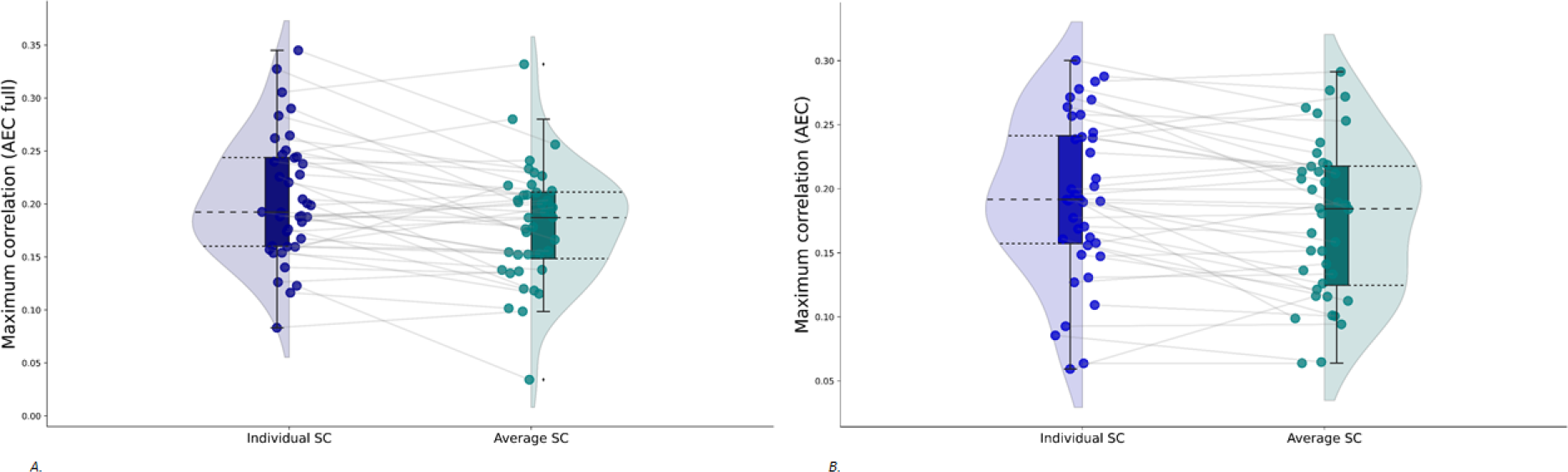
Paired rain cloud figures containing the maximum correlations, for all coupling values, between simulated and empirical FC. Grey lines between the dots connect one participant for simulations with the individual SC matrices as input to the model (blue rainclouds) and simulations with the group-averaged SC matrix as input to the model (green rainclouds). A. FC calculated with the AEC full. B. FC calculated with the AEC. Abbreviations: AEC: amplitude envelope correlation (AEC), AEC full: AEC computed over the full time-series.

We also computed a correlation between simulations with group-averaged SC and group-averaged FC, which showed a significant correlation between the two (AEC full: r = 0.40, *p* < 0.001 and AEC: r = 0.36, *p* < 0.001).

### Similarity between simulated versus empirical functional connectivity in non-matched versus matched participants

Next, we analyzed whether empirical FC of a given participant could be equally well predicted by simulated FC on the basis of another participant’s SC matrix. We correlated individually simulated FC to the empirical FC of all other participants. For both the AEC full and the AEC, in 5 out of the 40 participants, participants’ own individual correlation was significantly higher compared to the correlations with all other participants (see Figure 5).

**Figure 5.**
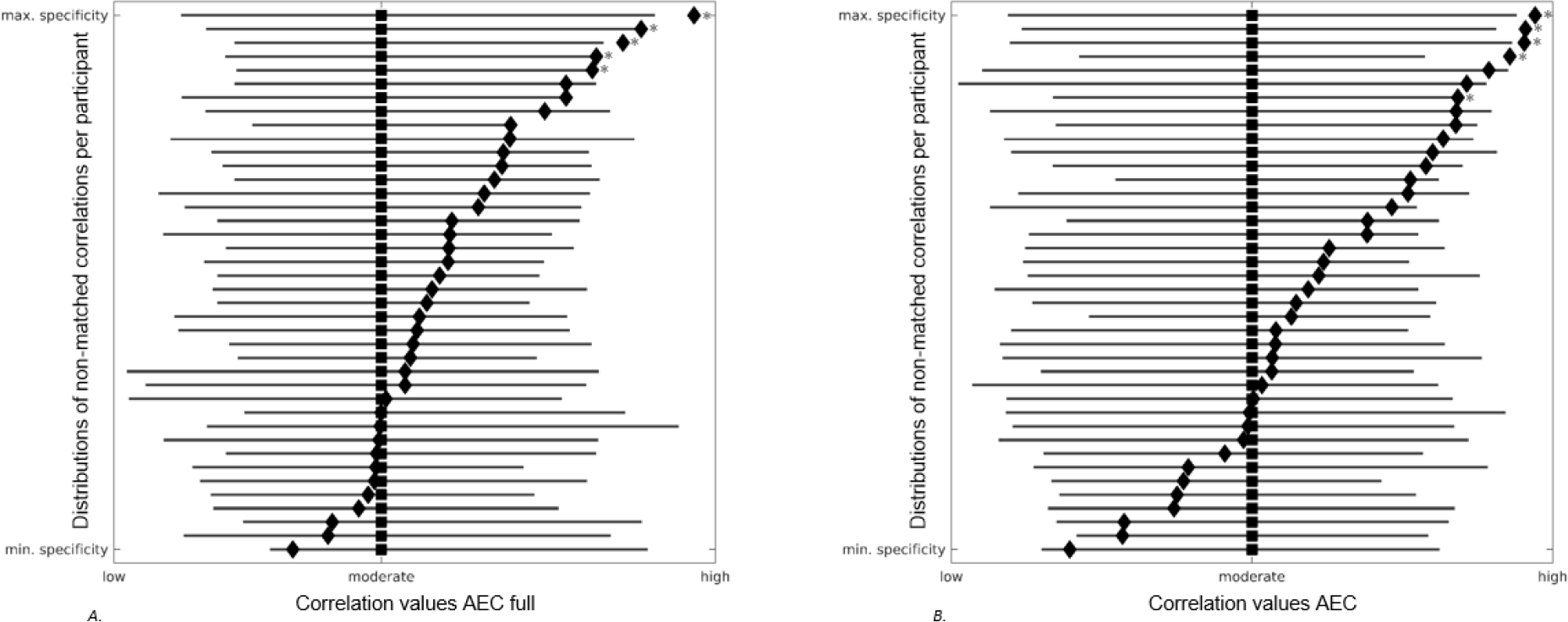
Forest plots showing the distributions of non-matched correlations per participant. Grey lines correspond to correlation distributions (low-moderate-high) between a participants own simulated FC and all other participants’ empirical FC. Black squares denote median values of these distributions. Black diamonds correspond to the correlation between a participants own simulated and empirical FC. Red stars display the correlations between participants own simulated and empirical FC that were significantly higher compared to correlations between participants own simulated FC and all other participants’ empirical FC. Participants are ranked based on the distance between their own correlation value and the median of all other correlation values, indicating the range between minimum and maximum specificity of participants own correlation values. A. AEC full. B. AEC. Abbreviations: AEC: amplitude envelope correlation (AEC), AEC full: AEC computed over the full time-series.

## Discussion

The main aim of this study was to assess the feasibility and accuracy of modeling individual empirical FC using individual empirical SC matrices. We found moderate correlations between simulated and empirical FC using the amplitude-based AEC, while the phase-based metrics (PLI and PLV) performed significantly worse. Using individual SC, instead of group-averaged SC, improved the correlation between simulated and individual empirical FC significantly. However, correlations between individually simulated FC and other participant’s empirical FC were in general not significantly lower than between the matched pair of FC patterns.

The FC simulations using individual SC outperformed simulations based on group-averaged SC, indicating increased precision modeling of brain activity and FC when incorporating participants’ own structural network. These findings are corroborated by Aerts and colleagues (Aerts et al 2018), who simulated fMRI data in brain tumor patients using The Virtual Brain. Individually optimized model parameters also resulted in improved accuracy of individually simulated FC. However, when correlating an individual’s simulated FC to the empirical FC of other participants, we found correlations that were comparable to matched simulated and empirical individual FC. Although this finding is in line with earlier work (Finger et al 2016), it remains unclear whether simulated FC can be attributed to a specific individual. It would be useful for future work to explore the causes of this apparent aspecificity. A recent study reported on subject specific MEG FC patterns, also known as functional fingerprints (Da Silva Castanheira et al 2021). Future studies could look into such fingerprints in repeated MEG measurements over time, both between and within participants. The variation that is present between and within participants in the match between simulated and empirical FC could increase our understanding of whether the simulated or empirical FC is underlying the aspecificity that we found.

A second main result of this study is the clear difference between amplitude- and phase-based metrics in the correlations between individually simulated and empirical FC. The AEC full and AEC outperformed the PLI and PLV, while the PLV performed better in comparison to the PLI. These findings partly corroborate earlier work in which only phase-based metrics were considered (Finger et al 2016), also showing better performance for the PLV in comparison to PLI. It is however important to note that Finger and colleagues used FC metrics both corrected and uncorrected for signal leakage. Although signal leakage is known to cause spurious correlations between nearby sources (Gross et al 2013), the previously mentioned study corrected their empirical data dependent on the FC metric. Since leakage is not present in our simulated data, we therefore chose not to perform leakage correction to our simulated data, but only to the empirical data. Important to note here however is that the PLI inherently corrects for leakage and therefore is corrected in both our simulated and empirical data. The difference in the performance of phase- and amplitude-based metrics could relate to the consistency levels of the FC metrics. In the alpha band, the AEC has been shown to be more consistent in repeated empirical measurements from the same participants, hypothetically since phase-based metrics are more susceptible to noise (Colclough et al 2016, Tewarie et al 2019b). If noise indeed underlies the poorer performance of phase-based FC metrics in individual simulations, including more data, i.e. including ten-instead of five-minute recordings, might improve results with these metrics (Liuzzi et al 2017). Additionally, previous research including EEG data of patients with Alzheimer’s disease found higher reproducibility of the PLI in the theta band, while the AEC was more consistent in the alpha and beta frequency bands (Briels et al 2020). This work could indicate that consistency of FC metrics might be frequency-dependent in empirical data, an aspect that we did not take into account by only analyzing our data in the alpha frequency band.

We found moderate (r=0.19 on average) correlations between individually simulated FC and individual empirical FC, which is lower than obtained by Finger and colleagues (average correlation of 0.53). However, direct comparison of these correlation values is not straightforward due to the many methodological differences between their study and ours. Nevertheless, several factors may have contributed to these results. The quality of both the empirical SC and FC matrices could have influenced the correlation strengths that we found. Regarding SC, tractography is known to underestimate the presence of interhemispheric fibers, which strongly influences modeling results (Messe et al 2015). The tractography method we used is the current standard in the field and takes care of false positives (Maier-Hein et al 2018). Nonetheless, future studies may investigate whether increasing the quality of the SC matrices, for instance by improving scanner hardware, diffusion sequences, duration of scans, or the tractography methods, could enhance modeling accuracy. Furthermore, MEG data is known to be susceptible to noise caused by environmental, instrumental and biological factors. Although we only included MEG data that was visually free from artefacts, noise may still have been present in the individual FC matrices. In an additional analysis we only took the strongest connections of the simulated and empirical FC into account (Figure S2), thereby decreasing the noise of the included connections. For the AEC, the resulting match between simulated and empirical FC was higher compared to taking the full matrices into account. Furthermore, functional connections can also occur where there are few or no structural connections, possibly explained by indirect connections and interregional distance (Meier et al 2016, Robinson 2012). This means that even small variations in SC can support many different FC patterns, which makes the interdependence between them complicated (Popovych et al 2018). Additionally, by correcting the empirical data for signal leakage, true zero-lag interactions are also removed which might have been present in the simulated data, causing a decrease in agreement between simulated and empirical data.

Computational models that use average SC as an input have been frequently applied so far (Abeysuriya et al 2018, Cabral et al 2014, Deco et al 2017, Hadida et al 2018, Moon et al 2015, O’Neill et al 2018, Tewarie et al 2019a, Tewarie et al 2014), but hamper further tailoring of such models to individuals, particularly in the setting of neurological disease modeling. Previously, damage that reflects different diseases, has been modelled with advanced computational models (Aerts et al 2020, de Haan et al 2012, Tewarie et al 2018, van Dellen et al 2013), but these disease models have not yet been applied to individual data. Such tailored disease models could elucidate mechanisms underlying functional network trajectories (Douw et al 2019) in neurological disease, for instance modeling the impact of focal lesions on global network dysfunction and cognitive decline.

To conclude, we show that simulated FC best relates to individual empirical FC when using the individual SC as input to the model, compared to the use of group-averaged SC. This work therefore underlines a first step towards individual FC modeling.

## Abbreviations

FC: functional connectivity
SC: structural connectivity
MEG: magnetoencephalography
PLI: phase lag index
PLV: phase locking value
AEC: amplitude envelope correlation
EEG: electroencephalography
MRI: magnetic resonance imaging
dMRI: diffusion MRI
FOD: fiber orientation distribution
AAL: anatomical labeling atlas

## Acknowledgements

The authors would like to thank Lucas Breedt for his help with the creation of the raincloud figures.

## Authorship confirmation statement

MDS, AH, MMS and PT had a major role in the acquisition of the data. SDK, LD, MMS and PT designed and conceptualized the study, did the analysis and interpretation of the data. SDK drafted the manuscript for intellectual and technical content. All other authors (LD, EvD, MDS, JJGG, CJS, AH, MMS, PT) also took part in writing, reviewing and revising the manuscript.

