## Supplementary information for "Modeling of individual neurophysiological brain connectivity"

**Supplementary Information manuscript *Modeling of individual neurophysiological brain connectivity***

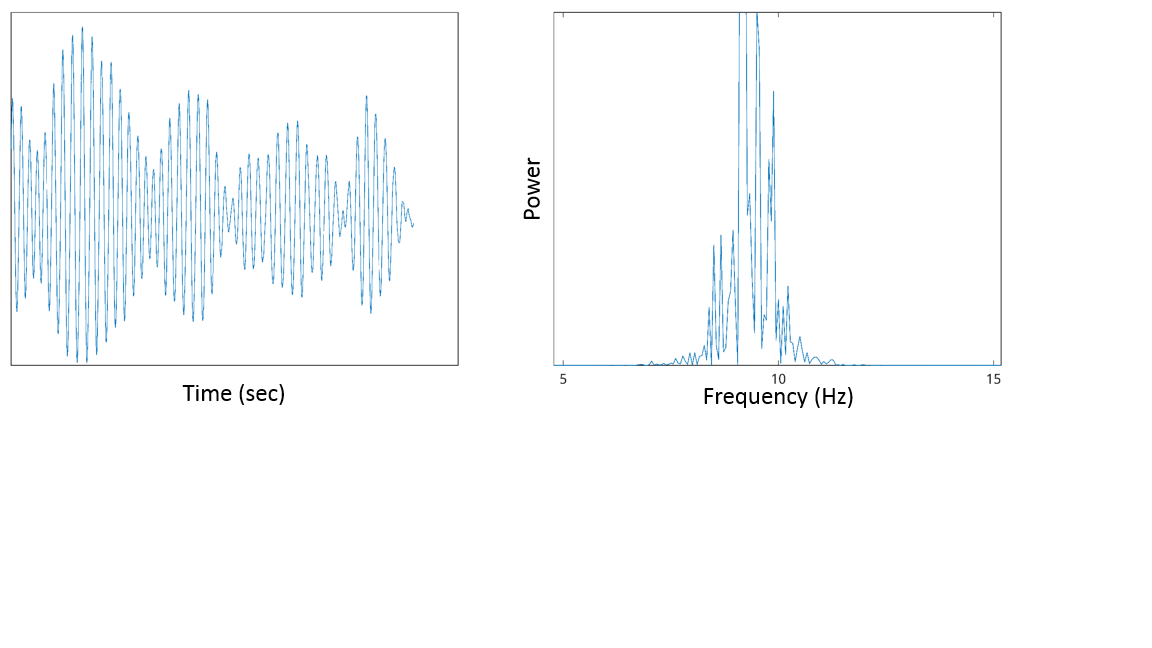

*Figure S1. Exemplar time-series (left) and power spectrum (right) of simulated data for one participant.*

*Table S1. Optimal coupling values per participant per FC metric*

| PARTICIPANT | Optimal coupling value | | | |
| --- | --- | --- | --- | --- |
|  | ***AEC full*** | ***AEC*** | ***PLI*** | ***PLV*** |
| 1 | 0.208 | 0.160 | 0.124 | 0.196 |
| 2 | 0.232 | 0.160 | 0.112 | 0.232 |
| 3 | 0.256 | 0.244 | 0.220 | 0.232 |
| 4 | 0.184 | 0.184 | 0.244 | 0.292 |
| 5 | 0.196 | 0.160 | 0.256 | 0.292 |
| 6 | 0.184 | 0.232 | 0.100 | 0.100 |
| 7 | 0.196 | 0.196 | 0.184 | 0.292 |
| 8 | 0.356* | 0.208 | 0.172 | 0.184 |
| 9 | 0.220 | 0.172 | 0.124 | 0.172 |
| 10 | 0.160 | 0.100 | 0.148 | 0.184 |
| 11 | 0.136 | 0.136 | 0.220 | 0.160 |
| 12 | 0.244 | 0.244 | 0.160 | 0.220 |
| 13 | 0.172 | 0.124 | 0.136 | 0.308* |
| 14 | 0.208 | 0.208 | 0.232 | 0.184 |
| 15 | 0.160 | 0.184 | 0.136 | 0.136 |
| 16 | 0.148 | 0.220 | 0.220 | 0.184 |
| 17 | 0.244 | 0.196 | 0.124 | 0.172 |
| 18 | 0.368* | 0.356* | 0.112 | 0.196 |
| 19 | 0.100 | 0.208 | 0.244 | 0.244 |
| 20 | 0.208 | 0.136 | 0.232 | 0.136 |
| 21 | 0.256 | 0.256 | 0.136 | 0.184 |
| 22 | 0.160 | 0.148 | 0.208 | 0.208 |
| 23 | 0.172 | 0.196 | 0.292 | 0.244 |
| 24 | 0.172 | 0.172 | 0.148 | 0.100 |
| 25 | 0.292 | 0.112 | 0.184 | 0.148 |
| 26 | 0.160 | 0.148 | 0.196 | 0.196 |
| 27 | 0.196 | 0.112 | 0.232 | 0.124 |
| 28 | 0.292 | 0.100 | 0.112 | 0.112 |
| 29 | 0.220 | 0.220 | 0.148 | 0.100 |
| 30 | 0.160 | 0.160 | 0.232 | 0.232 |
| 31 | 0.148 | 0.220 | 0.112 | 0.160 |
| 32 | 0.100 | 0.100 | 0.172 | 0.112 |
| 33 | 0.196 | 0.136 | 0.100 | 0.292 |
| 34 | 0.232 | 0.232 | 0.220 | 0.256 |
| 35 | 0.184 | 0.256 | 0.196 | 0.172 |
| 36 | 0.148 | 0.172 | 0.100 | 0.208 |
| 37 | 0.232 | 0.160 | 0.232 | 0.208 |
| 38 | 0.208 | 0.124 | 0.160 | 0.100 |
| 39 | 0.232 | 0.232 | 0.148 | 0.100 |
| 40 | 0.244 | 0.148 | 0.148 | 0.172 |

** Optimal coupling value fell within the higher coupling range: [0.1, 0.4]*

*Similarity between the strongest connections of individually simulated and empirical functional connectivity*

In addition to the correlations between the individually simulated and empirical FC matrices, we also studied the match between the strongest connections of the individually simulated and empirical data. Therefore, we applied the efficiency cost optimization method by De Vico Fallani et al. (De Vico Fallani et al 2017), which selects the strongest connections in a matrix by setting a threshold that optimizes the trade-off between the efficiency and wiring cost of a network. With this method, the strongest connections in the individual empirical data were determined and subsequently correlated to the values of the same connections in the individually simulated data. Per participant, the maximal correlation was determined as described in section ‘*Similarity between simulated and participant-specific empirical functional connectivity using individual structural connectivity’.* Differences between maximal correlations based on the full simulated and empirical matrices and the maximal correlations based on the strongest connections were tested with a Wilcoxon signed rank test.

For the AEC, maximal correlations between the strongest connections of simulated and empirical data showed to be significantly higher compared to the maximal correlations when the full matrices were taken into account (W = 205, *p* = 0.006, see Figure S2). For the AEC full, no differences were found between the maximal correlations based on the strongest connections and the maximal correlations based on the full matrices (W =453, *p* = 0.56).

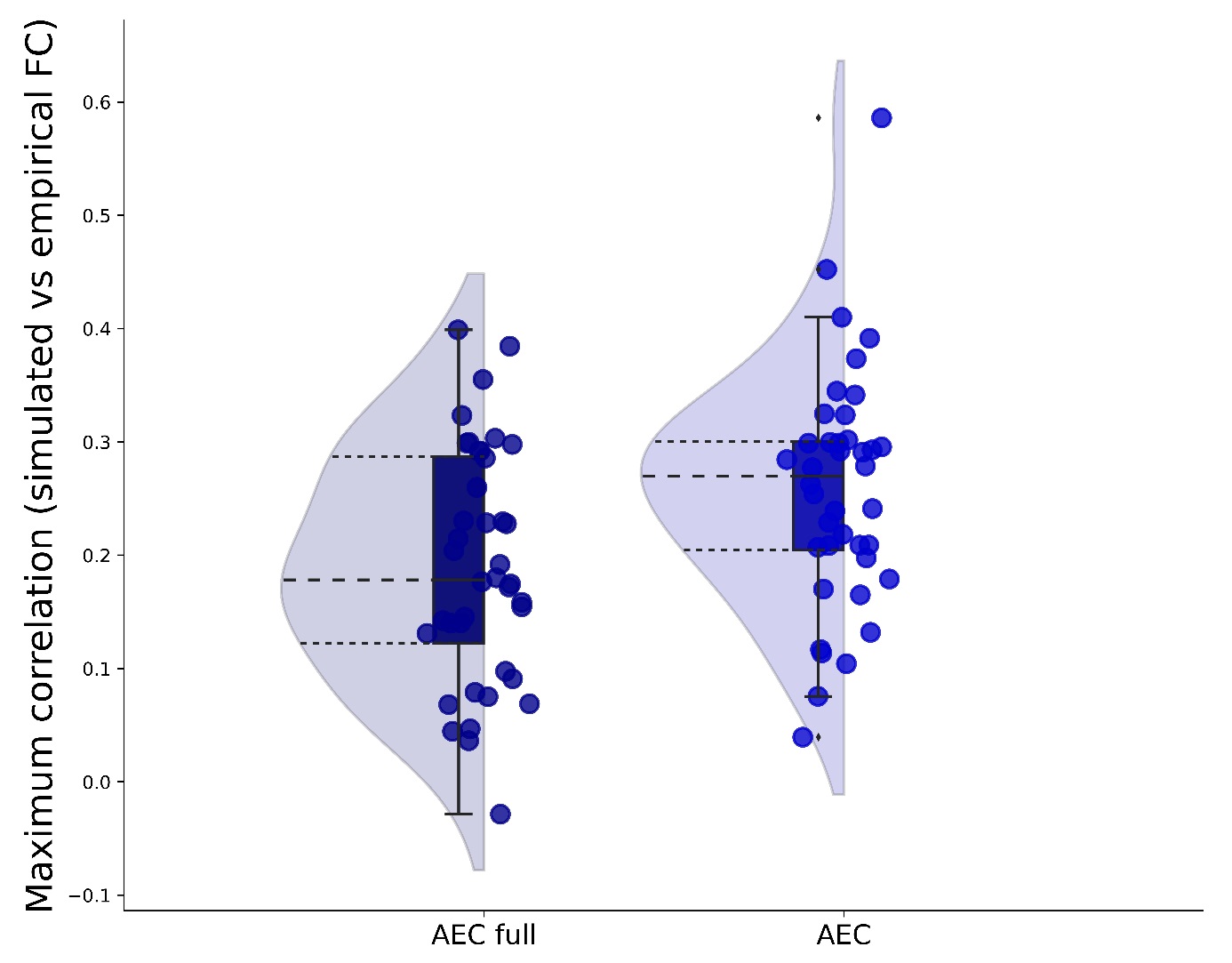

*Figure S2. Maximum correlations between the strongest connections of simulated and empirical FC.
Raincloud figures showing the maximum correlations between the strongest connections of simulated and empirical FC. A. FC calculated with the AEC full. B. FC calculated with the AEC. Abbreviations: AEC: amplitude envelope correlation (AEC), AEC full: AEC computed over the full time-series.*
